# How clinically relevant are prostate cancer cell lines? A comprehensive characterisation and multiomics comparison

**DOI:** 10.1101/2024.03.20.585982

**Authors:** Zahra Ahmed, Warda Mosabbir, Devansh Tandon, Snehal Pinto Pereira, Umber Cheema, Marilena Loizidou, John Withington, Caroline Moore, Uzoamaka Okoli, Susan Heavey

## Abstract

Cell line experiments arguably remain the most used tool in preclinical cancer research, despite their limitations. With almost 95% drugs entering human trials failing, and up to 90% preclinical research failing before even being tested in humans, we must shift the pre-clinical paradigm. A range of in silico, in vitro, in vivo and ex vivo approaches are gaining popularity, with the aim of potentially replacing cell line use. However, we cannot ignore the plethora of historical data from cell lines, nor write off their future use– especially within advanced bioengineered models. Therefore, we must question if and how cell lines hold clinical relevance. This study evaluates the clinical characteristics of 46 prostate cancer cell lines against worldwide data and investigates the biological features of seven cell lines in depth, comparing them to over 10,000 well characterised human cases from 24 studies in nine countries. Clinical features compared included age, ethnicity, Gleason grade, cancer type, treatment history and multiomics variables included mutations, copy number alterations, structural variants, microsatellite instability, mRNA and protein expression, and tumour mutational burden. We found that the most used cell lines accurately represent a minute proportion of prostate cancer patients. Furthermore, we recommend a pipeline for tailoring selection of clinically relevant cell lines with the ultimate aim of increasing the scientific methodology behind choosing a cell line.

## 1. Introduction

In drug development, an estimated 0.1% drugs which begin in preclinical trials are eventually approved (1) and oncology drugs have the highest attrition in clinical trials(2), due to incomplete proof of concept with human results failing to confirm preclinical findings (3, 4). The need to bridge the preclinical-clinical boundary has been stressed by the Academy of Medical Sciences and the Association of the British Pharmaceutical Industry (ABPI)(5), with proposed solutions including exploration of new preclinical and experimental models (6, 7).

Cell lines are frequently used for basic science, diagnostic and drug development; despite their shortcomings. The traditional use of cell lines in 2D demonstrates no heterogeneity, tumour microenvironment (TME) or immune influence (8). Furthermore, there is often cross-contamination (9). Despite this, cell lines have the advantage of not requiring biobank tissue or cell number restrictions.

Different PCa cell line models exist which are each derived from one patient and subsequently biologically altered during immortalisation (Supplementary Table 1).

Publicly available clinical datasets of PCa patients represent an opportunity to investigate the proportion of patients with specific PCa phenotypes and for analysis, visualization tools like cBioPortal encompass a range of clinical cohorts (Supplementary Table 2).

Limited PCa cell lines have been characterised using multiomics, but have not been compared comprehensively against patients, making them well characterised but not well validated. Important features to consider when assessing the clinical relevance of cell lines include donor ethnicity, age, cancer type, Gleason grade, metastatic stage, patient survival and androgen sensitivity.

A cell line cannot represent all patients but should hold clinical and biological relevance to the research question being asked. This study aimed to characterize and compare cell lines with publicly available patient data to determine how clinically and biologically relevant PCa cell lines and in what contexts. By tailoring the selection of cell line panels to match PCa target populations more accurately, we propose that a greater proportion of preclinical research can be utilised in practice.

## 2. Methods

### 2.1 Data collection, curation and comparison

Clinical and biological data was collected from a range of sources, overviewed in the Supplementary Methods and Supplementary Tables 3-6.

Seven cell lines from the 46 identified were analysed in depth using multiomics data from the CCLA Broad study and COSMIC. The most frequent mutations, CNAs and SNVs from the clinical cohort and cell lines were compared.

The seven cell lines from CCLE were compared against the KEGG PCa gene set (M13191,(10, 11, 12)) which acted as a control, and visual representations of gene interactions for altered genes were generated.

After identifying key gene alterations in the seven cell lines, the number of patients with any alteration the same was identified (Supplementary Table 7). Patients with increasing numbers of alterations the same were stratified, demonstrating increased biological similarity.

Using patient IDs with biological similarity to cell lines (Supplementary File 1), features of “cell line-like patients” were compared to one another, the overall clinical cohort and available global data. Based upon clinical and biological findings from this study, a guide to choosing relevant cell lines for preclinical research projects was composed.

### 2.3 Statistical analyses

Statistical analyses and graphs were plotted using GraphPad Prism and SPSS. Categorical variables were presented as frequencies and percentages, and continuous variables were checked for normality and compared using two way-ANOVA tests. For binary data, Chi Squared was used. Median survival was calculated using log-rank Mantel-Cox test and plotted on Kaplan-Meier curves. Significance was defined as p<0.05, alpha error 0.05.

## 3. Results

Collected clinical and biological data utilised in this study is shown in Figure 1.

**Figure 1.**
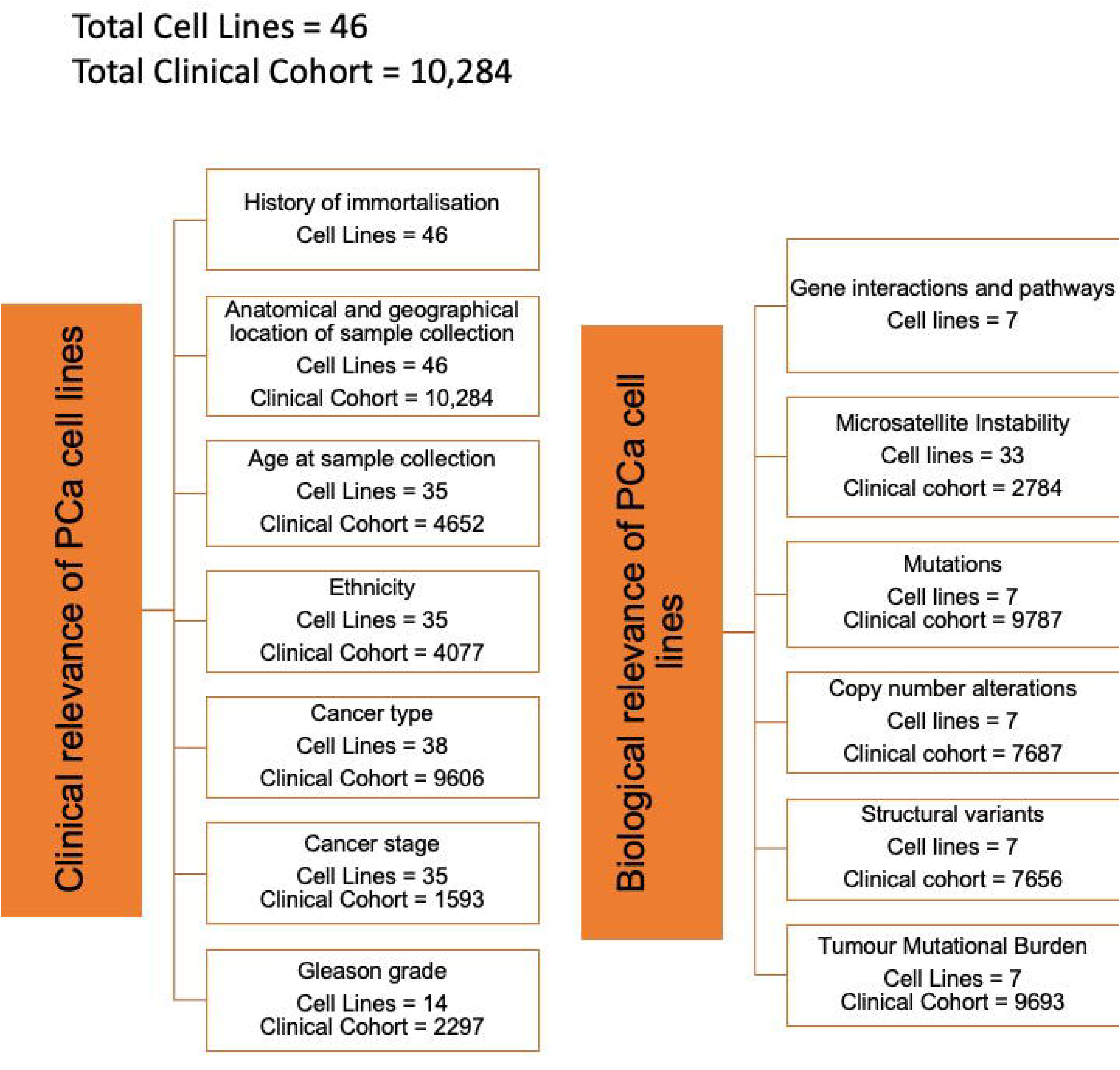
Overview of the clinical and biological factors were collected and analysed in this study. Raw data was collected for 7 major clinical factors. Raw data was collected for 6 overarching biological factors. The numbers of cell lines and size of clinical cohort for each factor is stated (if applicable).

**Figure 2.**
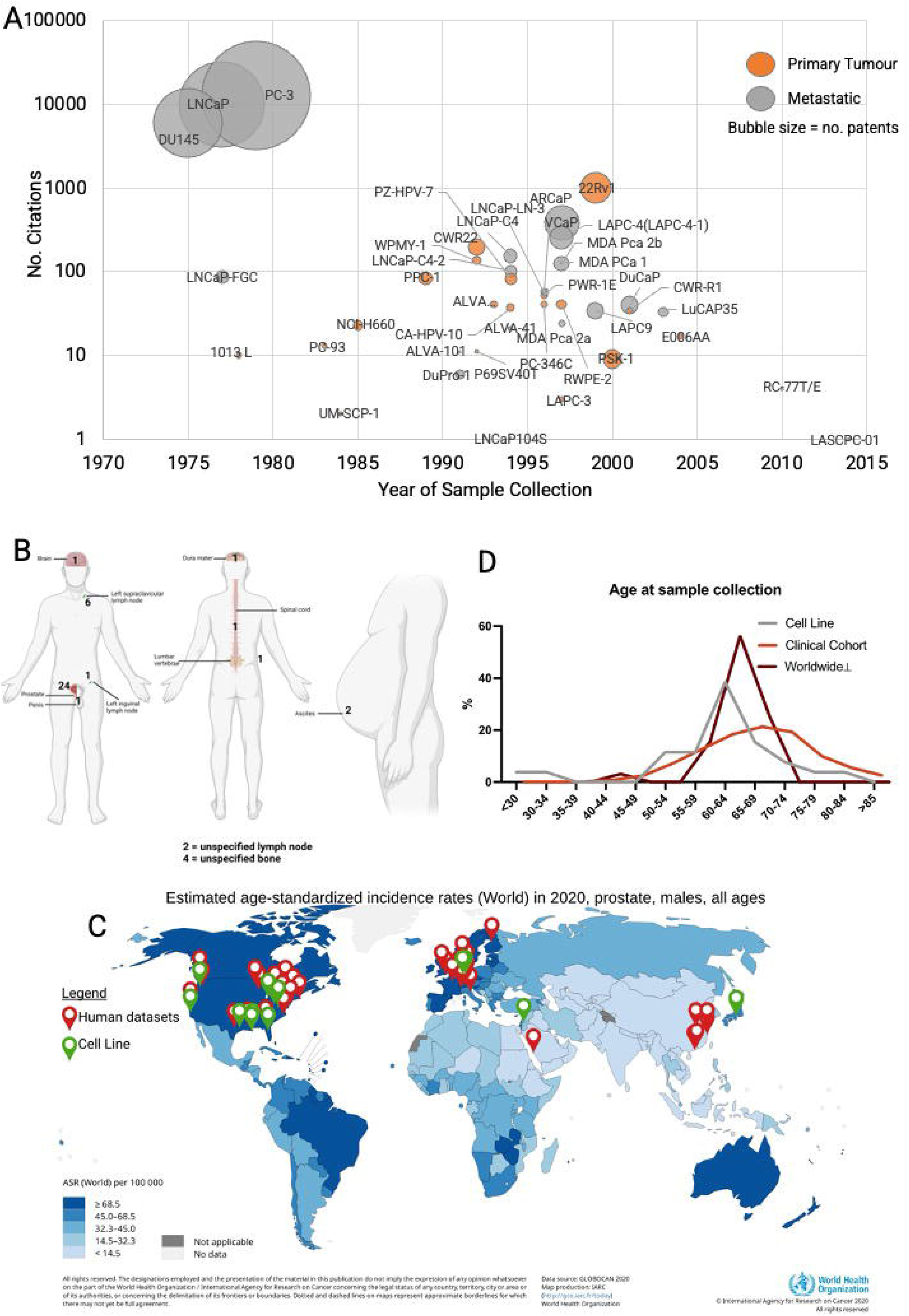
The most commonly used prostate cancer cell lines were derived over 40 years ago, were not derived from the prostate and are from very few geographical locations: **A** Bubble plot showing the most commonly used prostate cancer cell lines with number of literature citations (y axis) against the year of sample collection (x axis) with the size of bubble representing the number of patents and colour representing primary tumour (orange) or metastatic (grey) origin. **B** Illustration showing the anatomical locations of prostate cancer cell line sampling, numbers represent how many cell lines are derived from the anatomical site and the size of bubble represents the number of literature citations for cell lines derived from the anatomical site. **C** Map showing where cell line (green pins) and human datasets from publicly available clinical cohort data used in this study (red pins) were sampled from against a background gradient of prostate cancer incidence from low (light shade) to high incidence (dark shade). **D** Line graph showing distribution of age at sample collection for cell lines (N = 26), clinical cohort (N = 4646) and worldwide prostate cancer patients. ⊥ = Global Cancer Observatory, International Agency for Research on Cancer, World Health Organization, Cancer Today Website

### 3.1 Clinical relevance of prostate cancer cell lines

#### 3.1.1 History of immortalization

From a literature search, 46 PCa cell lines were identified and screened. PC-3 was the most cited and patent referenced, followed by LNCaP (Figure 3A). The three most used cell lines were immortalised over 40 years ago.

**Figure 3.**
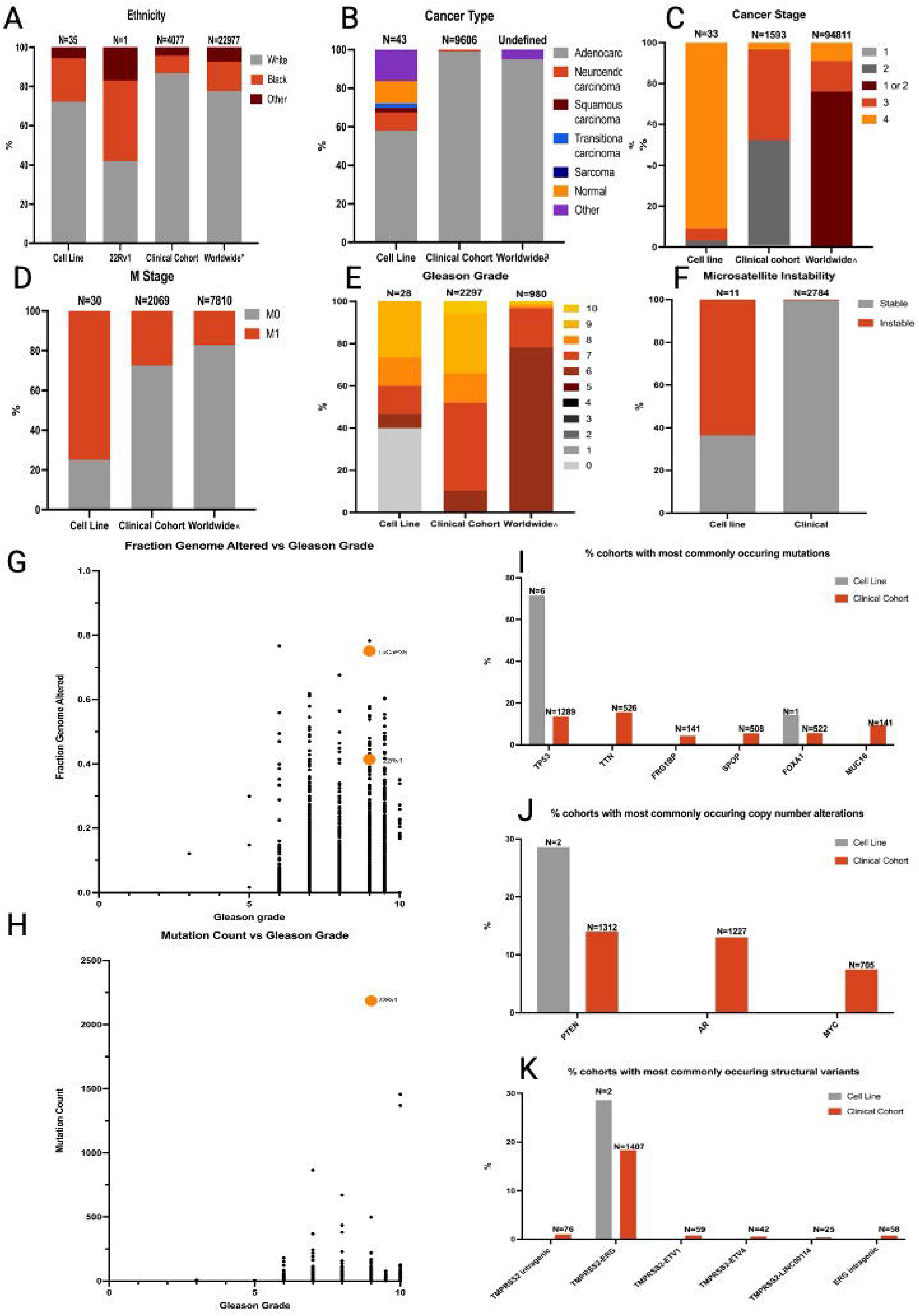
Cell lines and clinical cohorts have different clinical features to prostate cancer patients worldwide and the most common alterations in clinical datasets are not necessarily altered in cell lines: **A** Stacked bar chart showing the ethnicity distribution of original patients cell lines were derived from (N=35), 22Rv1 cell line, clinical cohort (N=4077) and worldwide prostate cancer patients (N=22977). Other races included Latino, Asian, Hawaiian, and Native American. **B** Stacked bar chart showing the cancer type of original patients cell lines were derived from (N=42), clinical cohort (N=9606) and worldwide prostate cancer patients. Other cancer types non-prostate carcinomas including bladder cancer. **C** Stacked bar chart showing the cancer stage of original patients cell lines were derived from (N=33), clinical cohort (N=1593) and worldwide prostate cancer patients (N=94811). **D** Stacked bar chart showing the proportion of patients with metastasis at sample collection of original patients cell lines were derived from (N=30), clinical cohort (N=2069) and worldwide prostate cancer patients (N=7810). **E** Stacked bar chart showing the distribution of Gleason grade of patients at sample collection of original patients cell lines were derived from (N=28), clinical cohort (N=2297) and worldwide prostate cancer patients (N=980). **F** Stacked bar chart showing proportion of patients with microsatellite instability for cell lines (N=11), clinical cohort (N=2784). **G** Scatter plot showing the fraction genome altered against Gleason grade for the clinical cohort and two cell lines with available data (LuCaP35 and 22Rv1). **H** Scatter plot showing the mutation count against Gleason grade for the clinical cohort and one cell line with available data (22Rv1). **I** The most common mutations out of 9787 patients from clinical datasets were TP53, TTN, FRG1BP, SPOP, FOXA1 and MUC16-mutations in these genes occurred in at least 10% patients respectively. Cell lines which also had mutations in TP53 were VCaP, DU145, 22Rv1, PC-3, LAPC4 and LAPC9. The cell line with a mutations in FOXA1 was DU145. **J** The most common copy number alterations (CNAs) out of 7656 patients from clinical datasets were PTEN, AR and MYC-CNAs in these genes occurred in at least 8% patients respectively. Cell lines with CNAs in PTEN were NCIH660 and LAPC9. **K** The most common structural variants (SVs) out of 7687 patients from clinical datasets were all fusions in either : TMPRSS2 (intragenic), TMPRSS2-ERG, TMPRSS2-ETV1, TMPRSS2-ETV4, TMPRSS2-LINC00114 and ERG (intragenic) - SVs in these genes occurred in at least 0.5% patients respectively. Cell lines also with fusions in TMPRSS2-ERG were NCIH660 and VCaP. M stage = Metastatic Stage, MSI = Microsatellite Instability. o = The Journal of the American Medical Association worldwide racial demographics of prostate cancer study and The Surveillance, Epidemiology, and End Results Program database, ∂ = Ackerman et.al Prostate surgical pathology, ∧ = Weerakoon et. al The current use of active surveillance in an Australian cohort of men

#### 3.1.2 Anatomical and geographical location of sample collection

Half PCa cell lines were from metastases (Supplementary Table 8, Figure 1B), most commonly from lymph nodes. 22Rv1 was the most used prostate-derived cell line.

Geographically, sample collection was concentrated mostly in Europe and North America. (Figure 1C).

#### 3.1.3 Age at sample collection

For PCa patients worldwide, age at sample collection most commonly occurred at age 60-64 (Figure 1D). For cell lines, almost 40% were also collected between 60-64, but the range was broader (29 to 83, Supplementary Table 8). The clinical cohort demonstrated an older average and normally distributed age (Supplementary Table 9).

#### 3.1.3 Ethnicity

Nomenclature for ethnicity was in line with guidance from the Office for National Statistics(13).Worldwide, Black patients accounted for 15.2% PCa diagnoses but only 9% of the total clinical cohort (Figure 3A). Seven cell lines were from Black patients and only two were from Asian patients (KuCaP13 and PSK-1). 22Rv1 comprised almost equal proportions of White and Black ancestry.

#### 3.1.4 Cancer type

Adenocarcinoma accounted for over 95% PCa types worldwide and 99.08% of the clinical cohort. Cell lines represented a range of cancer types (58.1% adenocarcinoma, Figure 3B).

#### 3.1.5 Cancer stage and metastatic stage

Over 90% cell lines were derived from patients with stage four PCa and 75% patients had known metastases (N=23). The clinical cohort and worldwide patients mostly had lower stage PCa (Figure 3C) without metastases (82.98% non-metastatic, Figure 3D).

#### 3.1.6 Gleason grade

Worldwide, 96.73% patients had Gleason grade 6 or 7. Almost half of the clinical cohort had Gleason grade >7 (48.06%, N=1103, Figure 3E). Gleason grade of donors from whom cell lines were established was only available for 15 cell lines and differed widely.

#### 3.1.7 Treatment History

Prior treatment history for the clinical cohort varied but was largely unknown and cell line donors received mostly no treatment or androgen deprivation therapy (Supplementary Table 10).

### 3.2 The biological relevance of prostate cancer cell lines

#### 3.2.1 Microsatellite Instability (MSI)

MSI was rare in the clinical cohort (0.43%) but more common in cell lines, with seven exhibiting MSI (63.64%, Figure 3F).

#### 3.2.2 Fraction genome altered and mutation count

Fraction genome altered, and mutation count, were plotted against Gleason grade. LuCaP35 was from Gleason 9 PCa and had a higher fraction genome altered compared to the clinical cohort with Gleason 9 (0.75 LuCaP35 vs median 0.12 clinical cohort, Figure 3G).

22Rv1 (also Gleason 9) had a mutation count 84 times higher than the median clinical cohort with Gleason 9 (N=298, Figure 3H).

#### 3.2.3 Mutations, Copy Number Alterations (CNAs) and Structural Variants (SVs)

The most frequent mutations, CNAs and SVs in the clinical cohort were compared against cell lines with multiomics data (Figure 3I-K). Cell lines typically lacked the most common CNAs and SVs from patients.

#### 3.2.4 Gene alterations and interactions

Multiomics data from cell lines was compared against the KEGG PCa gene set to evaluate how many genes were altered in each cell line (Figure 4A). LNCaP-FGC had the most alterations in genes from the KEGG PCa gene set, PC-3 had the least (37 and 10 respectively) and 24% genes were not altered in any cell line.

**Figure 4.**
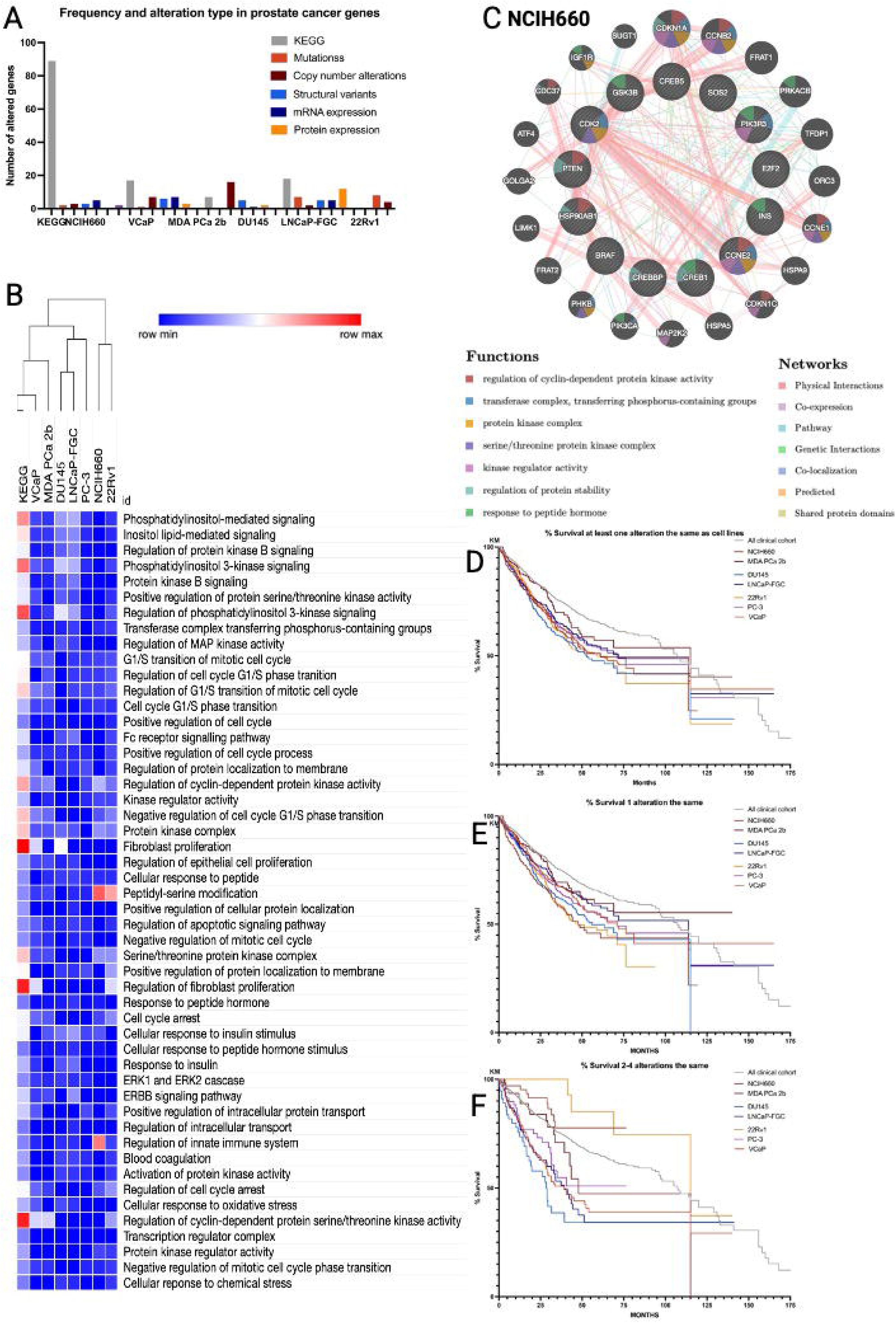
Cell lines do not share many of the same important gene alterations present in many prostate cancers; Comparing genes altered in cell lines at an individual and a pathway level: **A** Bar chart showing the frequency and alteration type in prostate cancer genes from the well characterised KEGG prostate cancer gene set were altered in. **B** Heatmap showing the proportion of genes altered from the KEGG prostate cancer gene set in each cell line/ total number of genes in the molecular pathway, where blue is lowest (0) and red is highest (0.26). Top 50 pathways with genes altered from KEGG prostate cancer gene set as a control on the left. **C** Identifying any gene interactions between altered genes in NCIH660 and their important predicted pathways which the cell line could be used to investigate. Diagram created using GeneMANIA with the inner ring represents genes altered in this cell line and the outer ring represents predicted related genes. The top seven pathways are colour coded inside the gene bubbles and the different coloured lines represent networks. **D-F** Kaplan-Meier survival plots for cell line-like patients compared to overall clinical cohort control (n=2323). **D**-Patients with at least one alteration the same as cell lines (overall), E-Patients with one alteration the same as cell lines, F-Patients with 2-4 alterations the same as cell line. MSI = Microsatellite Instability, KM = Kaplan-Meier

#### 3.2.5 Molecular pathways

Figure 4B details the top 50 pathways with the most alterations in genes from the KEGG PCa gene set compared with the number of genes altered in cell lines in these pathways. NCIH660 had a high proportion of genes altered in the peptidyl-serine modification and regulation of the innate immune system (21 and 20% genes, Figure 4C) ; 22Rv1 also had genes altered in the peptidyl-serine modification pathway (18%). Visualizations for remaining cell lines including gene interactions and pathways are shown in Supplementary Figures 1-5.

### 3.3 Identification and characterisation of cell line-like patients

#### 3.3.1 Sub-cohort identification

Identification of sub-cohorts of patients that were wholly representative of specific cell lines in terms of molecular alterations was largely unsuccessful. VCaP had the most patients with the same alterations present, 17 of which had 5-6 alterations the same out of the 89 genes in the KEGG PCa gene set. Other cell lines had less than 0.07% patients with 5-6 alterations the same (range 0 to 3).

#### 3.3.2 Survival

Patients with at least one alteration the same as a cell line had significantly lower percentage survival at 50 months compared to the overall cohort (*p*<0.001 log rank Mantel-Cox, Figure 4E&F). Most patients with 2-4 alterations the same as cell lines had a lower 50 month percentage survival compared to control, except 22Rv1 (Figure 4G).

### 3.2 Creation of cell line selection tool

Figure 5 demonstrates a three-step process guiding choice of PCa cell line. Step one considers key clinical factors relevant to almost all research questions, which should be used to shortlist cell lines of desired cancer stage, treatment status and androgen sensitivity. Step two narrows choice of cell line, by considering clinical factors including: cancer type, anatomical site of sample collection, patient age and race. Finally, step three caters for more specific factors.

## 4. Discussion

Traditional 2D monoculture cell lines are employed to investigate mechanisms and test drugs(14). In vitro PCa models have advanced to better emulate in vivo PCa by incorporating cells in various setups and 3D cell line-based models(15). Molecular characterization and limited detail of the clinical features of the patients PCa cell lines were derived from exists. However, there is limited literature evaluating the extent to which cell lines represent patients(16, 17) and no studies investigating which cell line is most suited for investigating patient subtypes. Therefore, this study aimed to contextualise PCa cell lines against large patient cohorts and worldwide data, and to characterise cohorts biologically similar to specified cell lines.

We found that many cell lines were established from metastasis and patients with rare presentations of PCa. From this alone, PCa cell lines are poorly clinically relevant. Additionally, there were no cell lines and patient datasets from countries with the highest PCa incidence. Black men often have aggressive PCa (18, 19, 20); With limited cell line models or patient data from Africa, it is likely that most PCa research excludes patients with the lowest survival. Furthermore, Black men are underrepresented in clinical trials attributable to factors relating to interaction with healthcare services, institutional racism and unconscious bias (21). Our results emphasise the need to implement strategies increasing representation of Black men in PCa research.

PCa cell lines often come from unique cancer types; PSK-1 came from a prostatic small-cell carcinoma patient (<2% PCas)(22) with Klinefelter syndrome (0.1-0.2% prevalence)(23). Whilst interesting clinically, these cell lines hold little relevance to understanding most PCas. Cell lines are biased to be derived from aggressive tumours, from their greater propensity to grow indefinitely. Although advanced stage cancers are of interest due to unmet clinical need, in future, more cell lines representative of early stage PCa are needed.

Seven cell lines profiled for multiomics data shared gene alterations from the KEGG PCa gene set, however, 24% genes were not altered in any cell line. This could mean that failed preclinical drug trials in these cell lines are due to lack of alterations in pathways, wherein the drug’s mechanism is targeted. Evaluation of cell line choice should be performed to ascertain whether the drug may work in a more suitable model. Most cell lines do not have detailed multiomics data available, which may bias researchers to select cell lines with more data available in relation to their genes of interest, rather than other potentially more relevant cell lines that lack data.

NCIH660 showed significant gene interactions between CDK1, CCNE2 and INS, perhaps suggesting it is a good option to investigate pathways involving these genes or for patients with alterations in these genes.

PC-3 was the least biologically altered, perhaps suggesting it is representative of patients with less altered genomes and earlier stage PCa.

To identify patients biologically-like cell lines, a tenuous link was suggested: as the number of gene alterations the same increased, so did similarity. From the clinical cohort, VCaP had the most patients with gene alterations the same, potentially suggesting it represents more PCa patients. However, the key finding here was that similarity was lower than expected overall.

Many cell lines had high MSI(MSI-H), which positively correlates with treatment response to immune checkpoint inhibitors in advanced PCa (24, 25). Furthermore, patients with high TMB have greater likelihood of immune activation(26) and response to immunotherapeutics. Therefore, cell lines with high MSI and TMB may be suited to investigate new checkpoint inhibitors.

The characteristics of patients biologically similar to 22Rv1 were different to other cell line-like patients (Supplementary File 2) with increased survival rates. 22Rv1 exhibits MSI (27) and patients with 2-4 alterations the same as 22Rv1 had the highest proportion of MSI-H and high TMB of all groups (10 and 9.02% patients). This could suggest that 22Rv1 is most suitable for immunotherapeutic experiments.

The multitude of PCa cell lines available, and strikingly high attrition rate of pre-clinical PCa research, render it pivotal to follow a systematic, evidence-based approach when selecting cell lines for pre-clinical research. Figure 5 serves as a tool for this, and should be used to whatever degree deemed appropriate.

**Figure 5.**
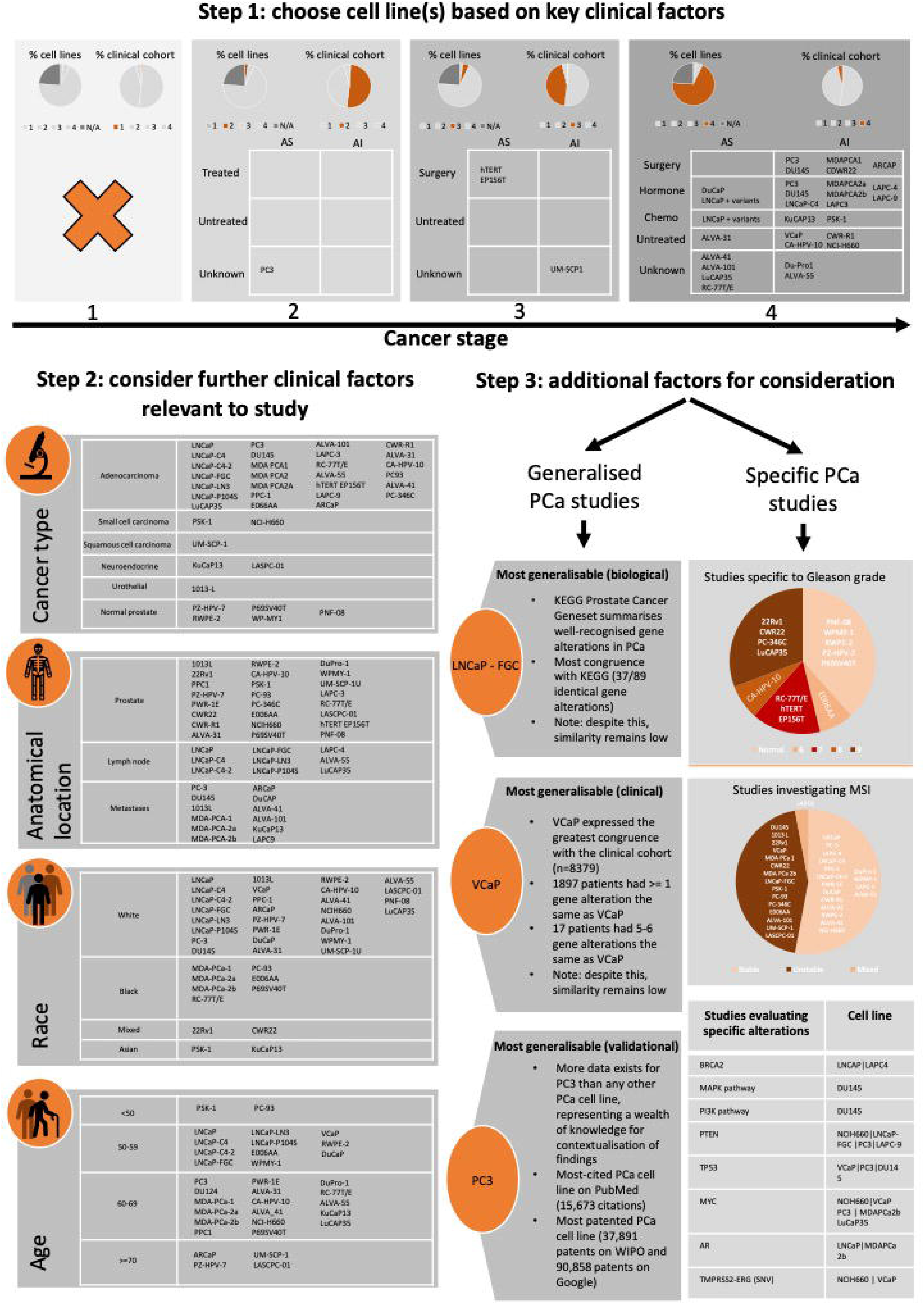
Selection of clinically relevant prostate cancer cell line(s) should be tailored to the research question. Researchers are encouraged to select a panel of cell lines to carry out experiments on. Each cell line can be selected based on a variety of factors, stratified into steps based on importance and impact. **A** Step 1: Cell lines can be selected based initially on their baseline clinical characteristics. Here, PCa cell lines are stratified based on cancer stage, the mode of treatment and whether the cancer is androgen sensitive or not. Pie charts highlight the discrepancy between the percentage of cell lines and percentage of the corresponding clinical cohort, based on donor characteristics at the time of data collection. **B** Step 2: Further clinical characteristics should be taken into consideration. Here, PCa cell lines are split into tables based on cancer type, anatomical location of sample collection, age of donor at time of sample collection and race of donor. **C** Step 3: Relevance to the research question should be used to further select cell lines for inclusion in a panel. Here, A flowchart categorises PCa cell lines based on whether the research question is generalised or specific in nature. The generalised side of the flowchart specifies 3 cell lines that may be suitable for predominantly broad studies. The specific side of the flowchart uses pie charts and tables to further stratify cell lines based on Gleason grade, microsatellite instability and specific gene alterations that are most popular in PCa research. A panel should include cell lines possessing an alteration of interest as well as cell lines that do not.

A study of this kind requires compromises to be made where global data is not available. Sources for Gleason grade and ethnicity were from single-country studies but included due to large patient cohorts. Another limitation of this study is that the number of cell line-like patients may have been skewed by alteration type and genes profiled for in each dataset. This was controlled for by including studies with a range of alterations (Supplementary Table 11).

Only seven PCa cell lines had multiomics data available, however, cell lines lacking multiomics data showed clinical and biological relevance. These cell lines should be comprehensively characterised to see if they hold relevance and if not, more representative cell lines are required.

Looking forward, 2D cell lines are likely to be continually used in 3D culture, especially as prostate-specific treatment studies have shown 3D in vitro models to be more biomimetic of in vivo tumours (28).

## 5. Conclusion

Cell lines are valuable in PCa research and are used for basic science research, development of advanced models, development of drugs and diagnostics, as well as training of junior scientists. However, it is pivotal that cell lines are applicable to the largest cohort of PCa patients possible and there are available niche cell lines representing patient subtypes. This study contextualised commonly used PCa cell lines against patients in clinical cohorts and worldwide. Cell line-like patients were characterised to inform researchers which cell line is most suited to their targeted cohort and experiment.

## Acknowledgments

Thank you to the Wolfson Foundation and the Royal College of Physicians for funding this project.

---

**Supplementary Table 1:**
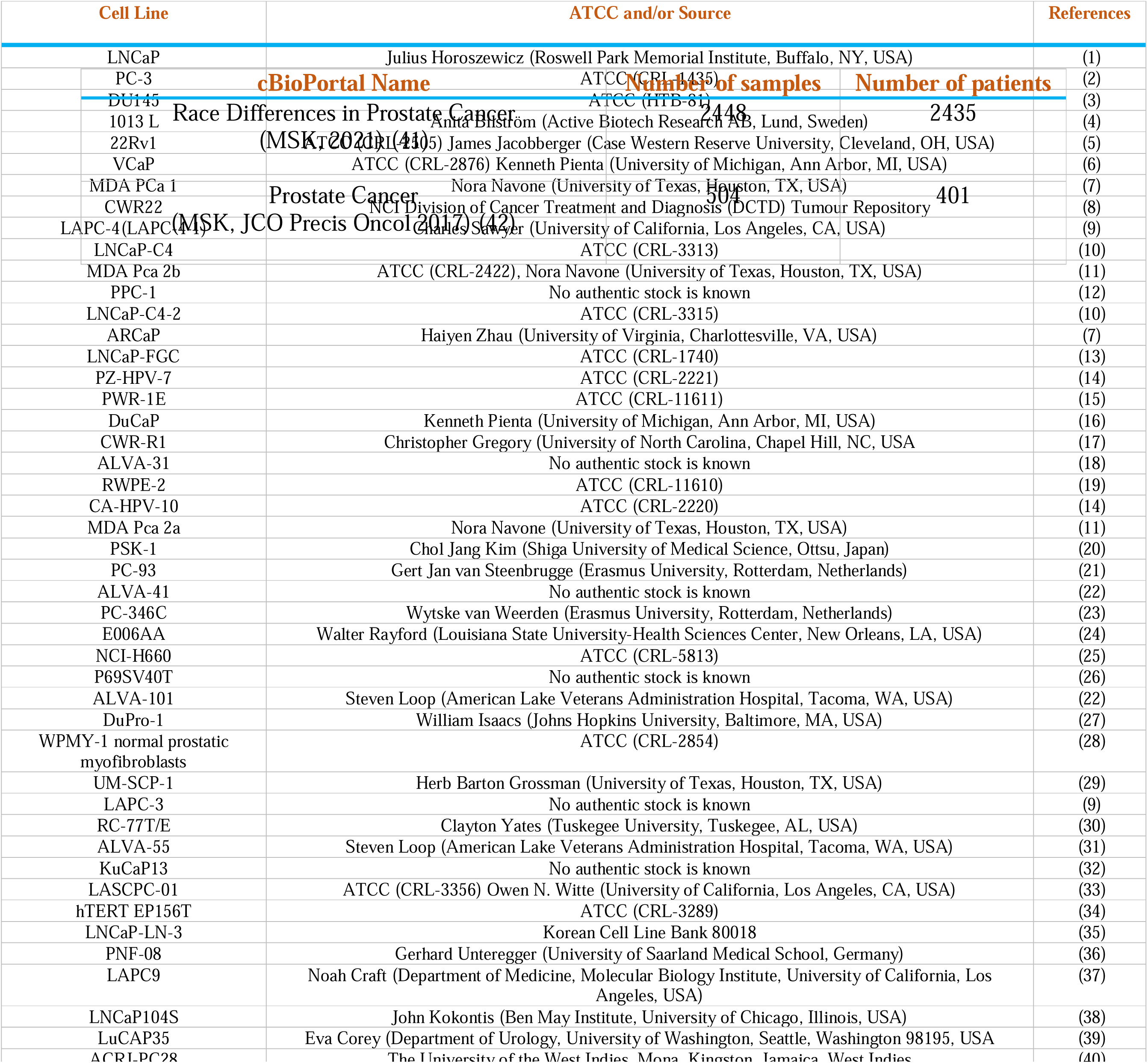
Prostate cancer cell lines analysed in this study.<colcnt=1>

**Supplementary Table 2:**
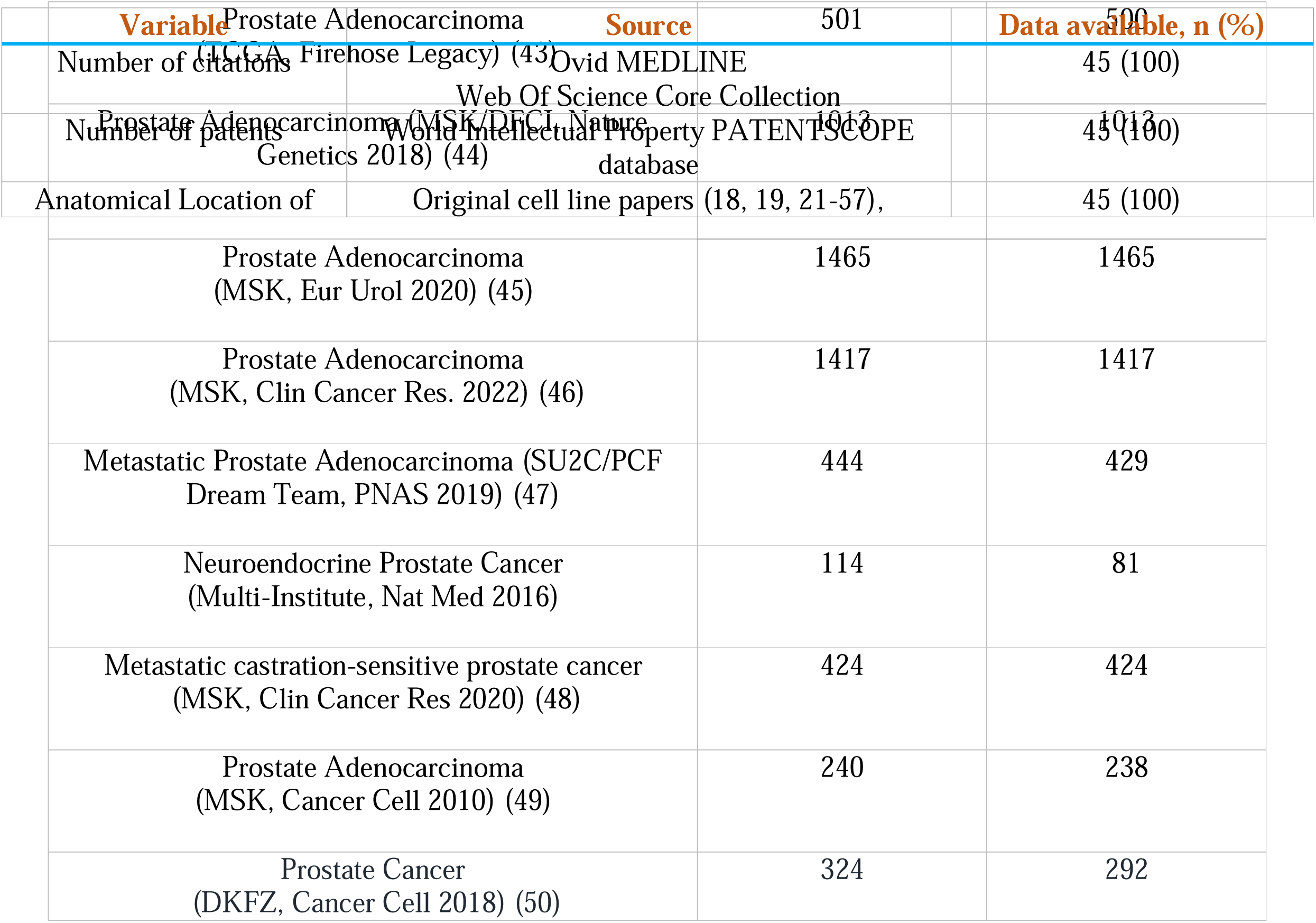
Clinical cohorts accessed on cBioPortal used in this study.

**Supplementary Table 3:** Baseline characteristics investigated for each cell line, COSMIC = Catalogue of Somatic Mutations in Cancer

|  |  |  |
| --- | --- | --- |
| sample collection | Cellosaurus website, ATCC website, COSMIC database |  |
| Patient age at sample collection | Original cell line papers (18, 19, 21-57), Cellosaurus website, ATCC website, COSMIC database | 27 (60) |
| Geographical location of sample collection | Original cell line papers (18, 19, 21-57) | 45 (100) |

**Supplementary Table 4 :**
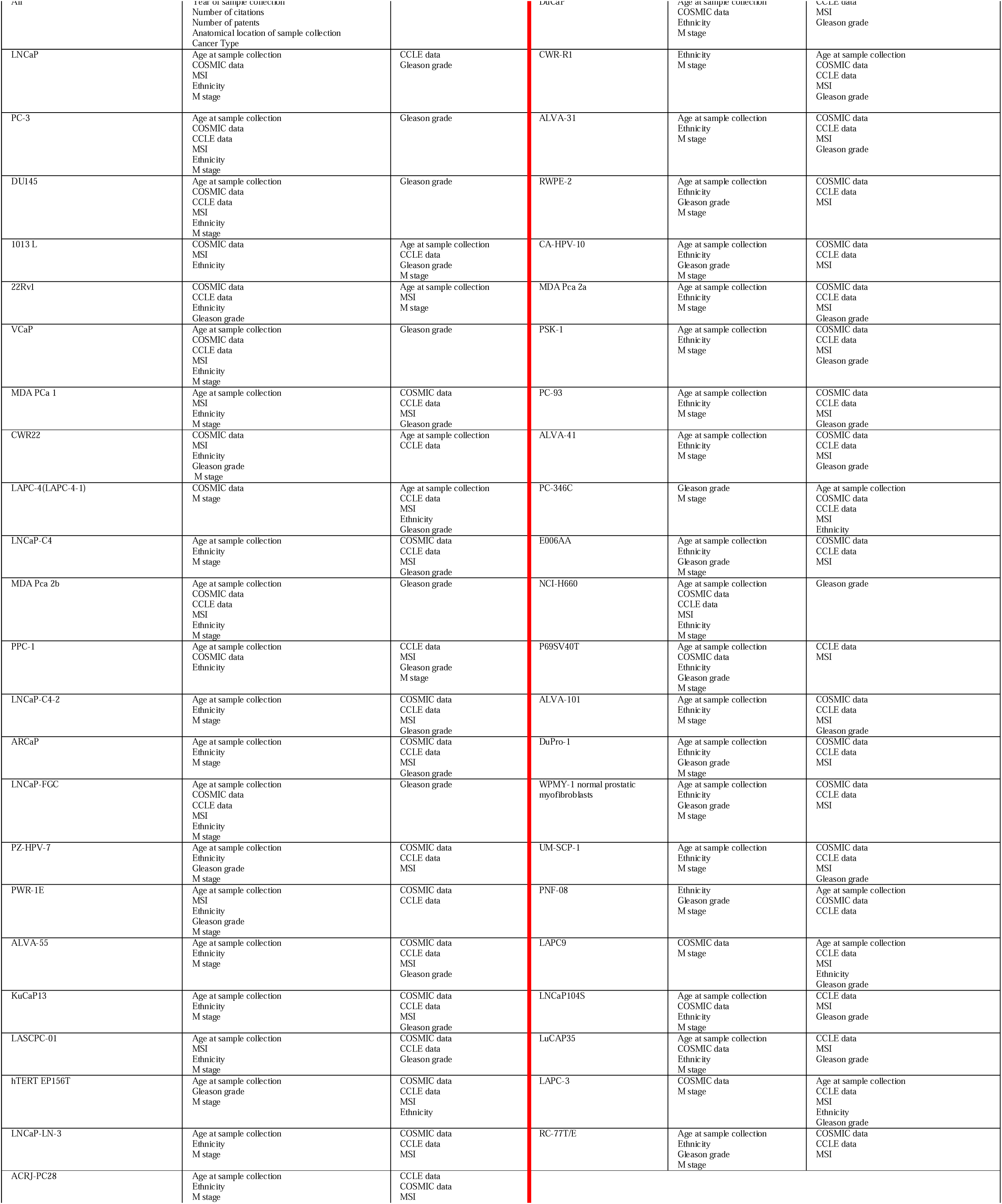
Data available for each cell line

**Supplementary Table 5:** Sources of worldwide Prostate Cancer data. GCO = Global Cancer Observatory, IARC = International Agency for Research on Cancer, WHO = World Health Organization. JAMA = The Journal of the American Medical Association, SEER = The Surveillance, Epidemiology, and End Results Program.

| Variable | Source |
| --- | --- |
| Patient age at sample collection (radical prostatectomy) | Systematic Review on robotic-assisted radical prostatectomy (51) |
| Geographical distribution of prostate cancer (incidence) | GCO, IARC, WHO Cancer Today Website (52, 53, 54) |
| Ethnicity of prostate cancer patients | JAMA worldwide racial demographics of prostate cancer study and SEER database (55) |
| Gleason grade of prostate cancer patients | Weerakoon et. al The current use of active surveillance in an Australian cohort of men (56) |
| Prostate cancer type | Ackerman et.al Prostate surgical pathology (57) |

**Supplementary Table 6:**
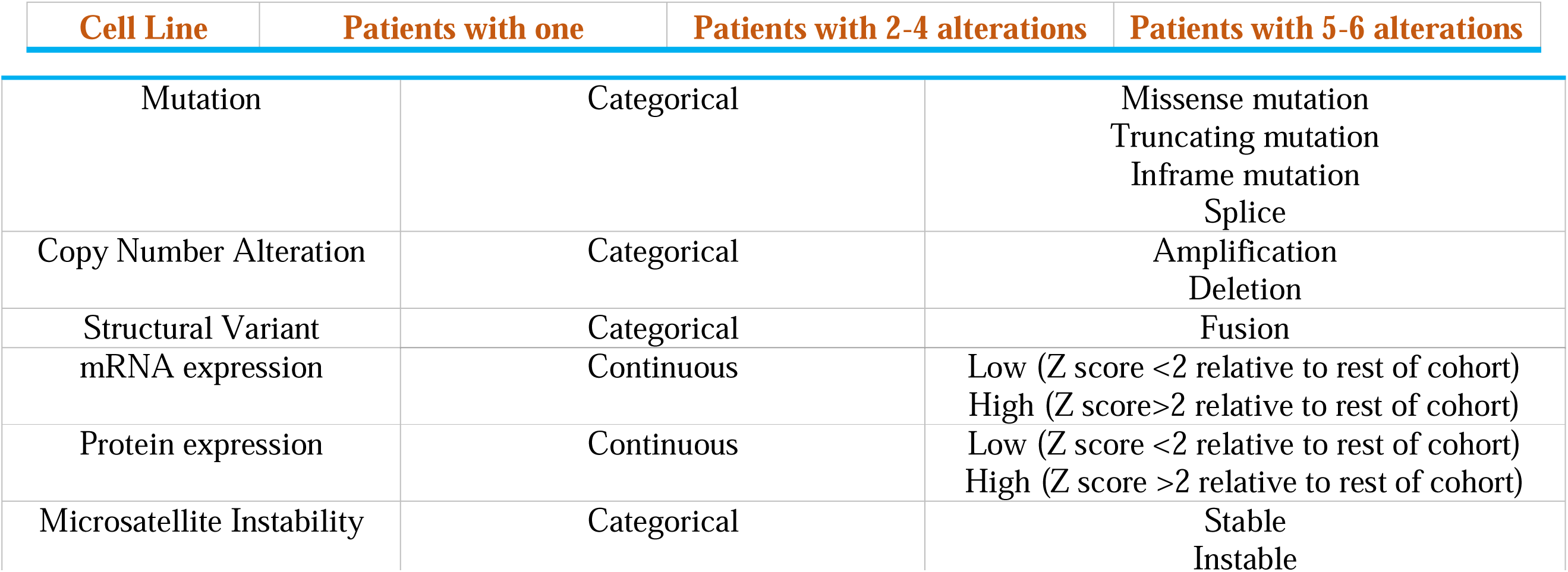
Biological variables investigated for each cell line.

| Cell Line | Patients with one | Patients with 2-4 alterations | Patients with 5-6 alterations |
| --- | --- | --- | --- |

|  |  |  |
| --- | --- | --- |
| Mutation | Categorical | Missense mutation<br>Truncating mutation<br>Inframe mutation<br>Splice |
| Copy Number Alteration | Categorical | Amplification<br>Deletion |
| Structural Variant | Categorical | Fusion |
| mRNA expression | Continuous | Low (Z score <2 relative to rest of cohort)<br>High (Z score >2 relative to rest of cohort) |
| Protein expression | Continuous | Low (Z score <2 relative to rest of cohort)<br>High (Z score >2 relative to rest of cohort) |
| Microsatellite Instability | Categorical | Stable<br>Unstable |

**Supplementary Table 7:**
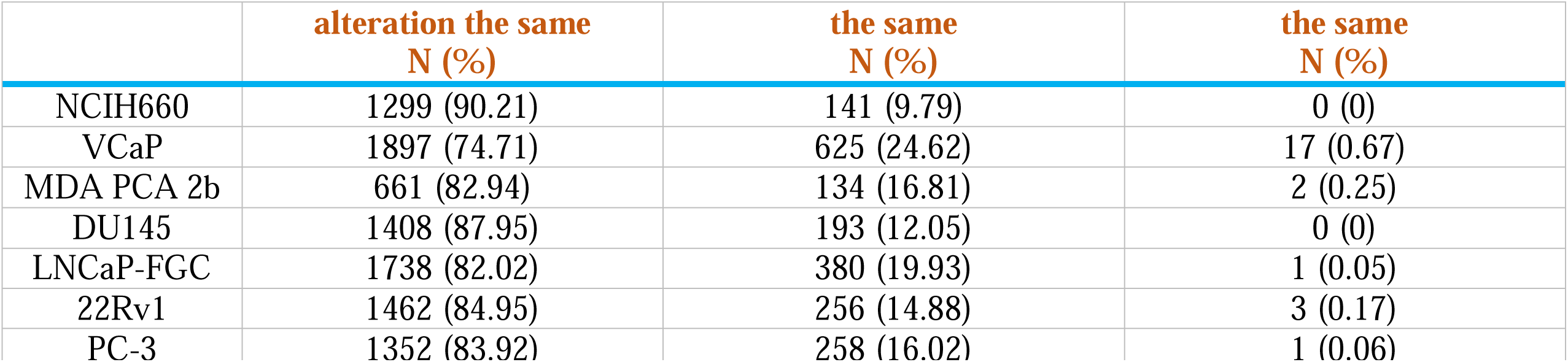
Number of patients with alterations the same as cell line

|  | **alteration the same**<br>**N (%)** | **the same**<br>**N (%)** | **the same**<br>**N (%)** |
| --- | --- | --- | --- |
| NCIH660 | 1299 (90.21) | 141 (9.79) | 0 (0) |
| VCaP | 1897 (74.71) | 625 (24.62) | 17 (0.67) |
| MDA PCA 2b | 661 (82.94) | 134 (16.81) | 2 (0.25) |
| DU145 | 1408 (87.95) | 193 (12.05) | 0 (0) |
| LNCaP-FGC | 1738 (82.02) | 380 (19.93) | 1 (0.05) |
| 22Rv1 | 1462 (84.95) | 256 (14.88) | 3 (0.17) |
| PC-3 | 1352 (83.92) | 258 (16.02) | 1 (0.06) |

**Supplementary Table 8:**
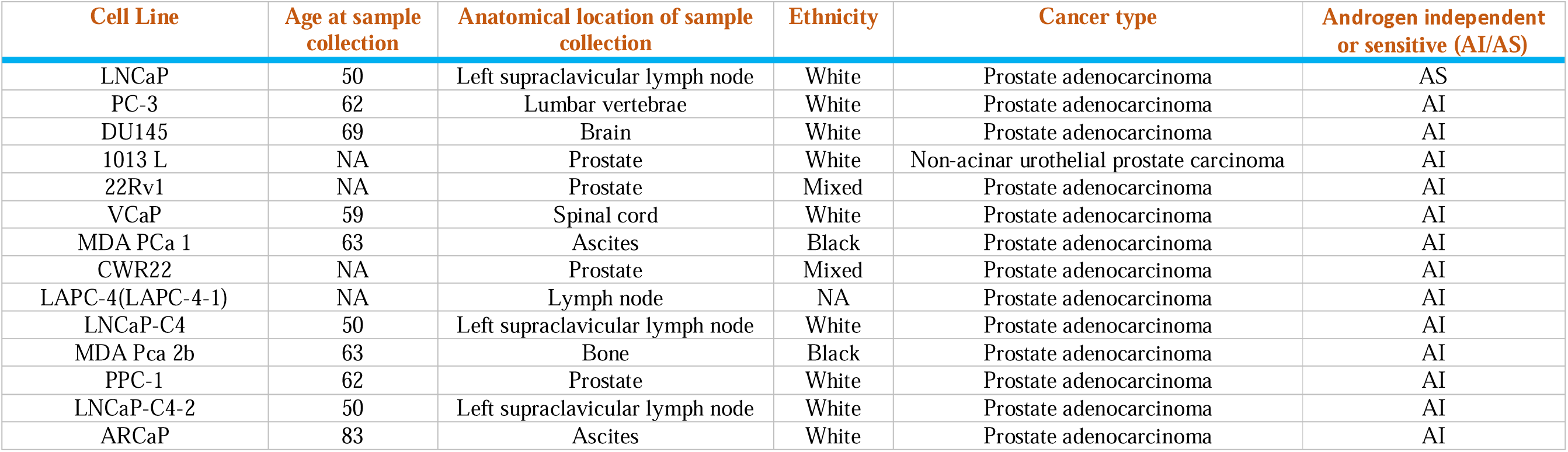
Key clinical variables for each cell line

**Supplementary Table 9:**
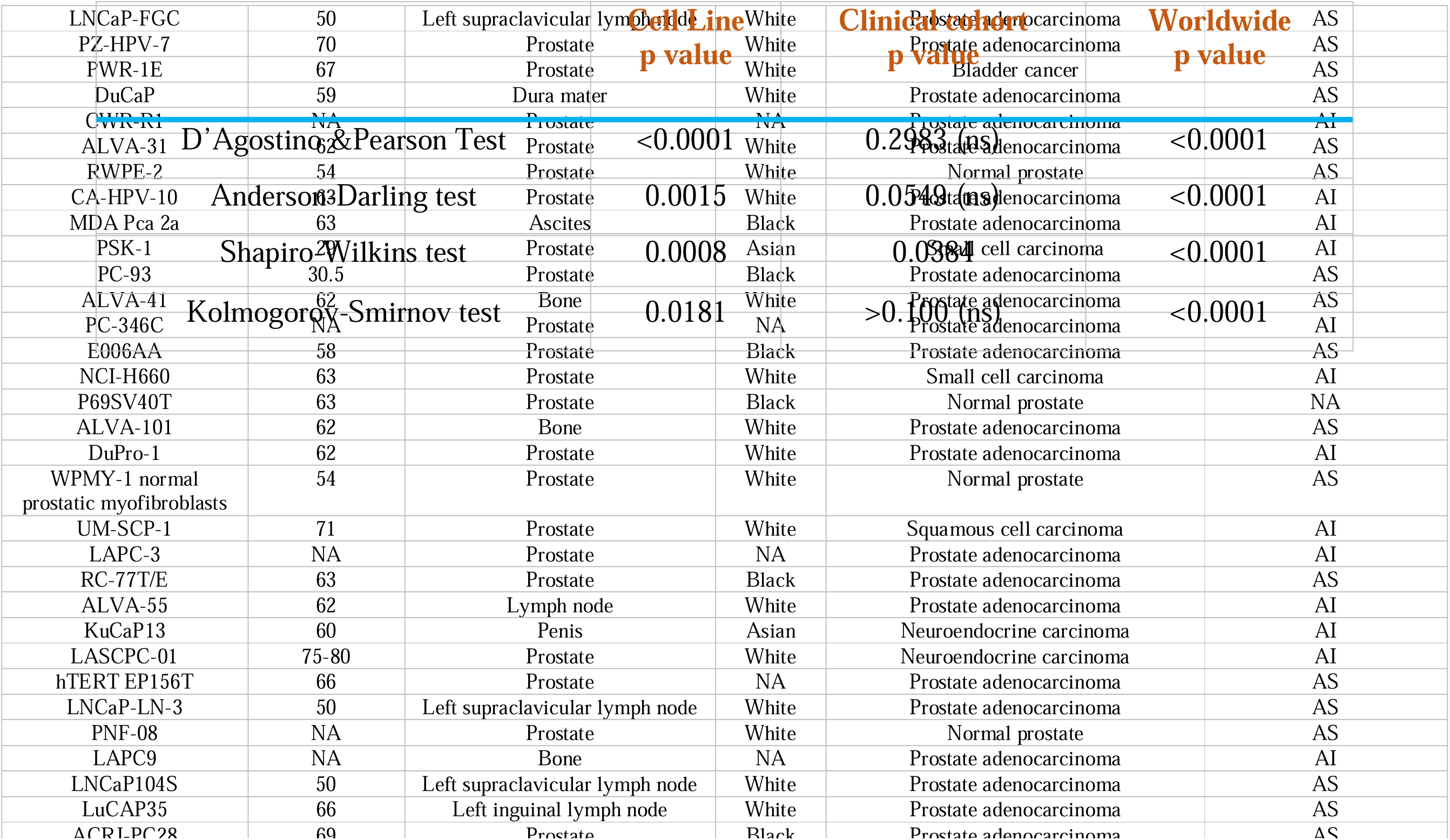
Results of normality distribution tests for distribution of age at sample collection.

**Supplementary Table 10:** Table demonstrating percentage of cell lines, or patients receiving different forms of treatment for prostate cancer prior to sampling

| **Treatment** | **Clinical Cohort (%)** | **Cell Line (%)** |
| --- | --- | --- |
| No treatment | 8.99 | 30.40 |
| ADT | 1.45 | 30.40 |
| Chemotherapy | 0.59 | 13.04 |
| Unknown | 88.97 | 28.26 |

**Supplementary Table 11:** Biological alterations screened for in clinical cohorts accessed on cBioPortal.

| cBioPortal Name | mRNA Expression | Protein Expression | Mutations | Copy Number Alterations | Structural Variants | Microsatellite Instability |
| --- | --- | --- | --- | --- | --- | --- |
| Race Differences in Prostate Cancer (MSK, 2021) (41) | ? |  | ? | ? | ? |  |
| Prostate Cancer (MSK, JCO Precis Oncol 2017) (42) |  |  | ? | ? | ? |  |
| Prostate Adenocarcinoma (TCGA, Firehose Legacy) (43) | ? | ? | ? | ? |  |  |
| Prostate Adenocarcinoma (MSK/DFCI, Nature Genetics 2018) (44) | ? | ? | ? | ? |  |  |
| Prostate Adenocarcinoma (MSK, Eur Urol 2020) (45) |  |  | ? | ? | ? | ? |
| Prostate Adenocarcinoma (MSK, Clin Cancer Res. 2022) (46) |  |  | ? | ? | ? | ? |
| Metastatic Prostate Adenocarcinoma (SU2C/PCF Dream Team, PNAS 2019) (47) | ? |  | ? | ? | ? |  |
| Neuroendocrine Prostate Cancer<br>(Multi-Institute, Nat Med 2016) | ? |  | ? |  |  |  |
| Metastatic castration-sensitive prostate<br>cancer<br>(MSK, Clin Cancer Res 2020) (48) |  |  | ? | ? | ? | ? |
| Prostate Adenocarcinoma<br>(MSK, Cancer Cell 2010) (49) | ? |  | ? | ? |  |  |
| Prostate Cancer<br>(DKFZ, Cancer Cell 2018) (50) | ? |  | ? |  |  |  |

